# Reaction diffusion plus growth: A numerical model for Turing morphogenesis and Ben-Jacob patterns

**DOI:** 10.1101/370304

**Authors:** Kai Trepka

## Abstract

Building models of organismal growth enables predictions of natural variability and responses to perturbations. Complex systems such as animal pattern development and bacterial colonies can be modeled numerically using a reaction-diffusion system with relatively few factors and yield qualitatively accurate results, allowing exploration of different equilibrium and non-equilibrium solutions. Here, a discrete numerical reaction diffusion system is used to qualitatively reproduce both common animal patterns and Ben-Jacob bacterial growth.

## INTRODUCTION

The diffusion equation has been used for decades to model processes from Brownian motion to population ecology. A particular focus has been placed on morphogenesis, or how a spherically symmetric egg becomes a bilateral organism. However, challenges arise when developing analytical solutions to coupled differential equations involving dozens of complicated biological factors and natural sources of randomness. Given the rise of computational power, numerically modeling diffusion poses a potential avenue for numerically modeling organismic growth and development.

Given a collection of small particles and a heterogeneous concentration profile, over time the particles will be pushed by random thermal fluctuations into a more uniform profile, yielding change in concentration that scales with the diffusion constant, D (Equation 1).

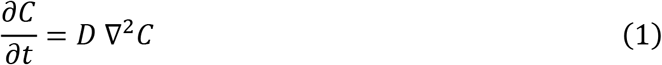

Generalizing this to two dimensions, the continuous concentration profile at a given timepoint is defined by Green’s function, G(x,t), with constant n representing the number of timesteps and t the increment per timestep (Equation 2).

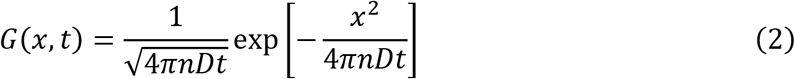

## METHODS

### The Diffusion Equation

A discretized, two-dimensional version of this model was implemented in Matlab using a matrix that keeps track of the location of each diffusing species at each timepoint, and updates subsequent timepoints based on the previous timepoint, the diffusion constant, and a discretized version of the Laplacian from the diffusion equation. This discretized Laplacian will take all the concentration from a given point, divide it into four parts (a two-dimensional grid square has four neighbors), and move the current concentrations “next door.” In other words, regions with many particles will “lose” their particles to surrounding regions with fewer particles.

### Reaction and Growth

Many physical phenomena are limited by diffusion, but are also influenced by the secretion or absorption of certain particles, independent of diffusion itself. In this case, the diffusion equation is modified for the change in concentration with respect to time to incorporate these outside factors.

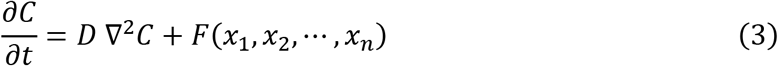

In general, there is an additive modification F that is some function of x_i_’s, the local concentrations of factors (Equation 3). In other words, we model the diffusion of factors away from a position, combined with processes that regenerate or absorb the product at discrete positions.

In addition to diffusing factors, larger, more static objects are relevant when considering biological systems. These objects can interact with diffusing factors in a variety of ways – secreting them, absorbing them, dividing or dying. Adding interactions is challenging analytically, but easy numerically. To do so, the Matlab model defines a set of equations for how objects interact with factors in solution, and keeps track the locations and concentrations of each item at every discretized timepoint.

### System Validation

As a basis for this model, a 2D Laplacian operator in Matlab was verified to produce the expected concentration profile for a diffusing point source. The Gaussian concentration profile after 1000 timesteps is shown (Figure 1).

**Figure 1:**
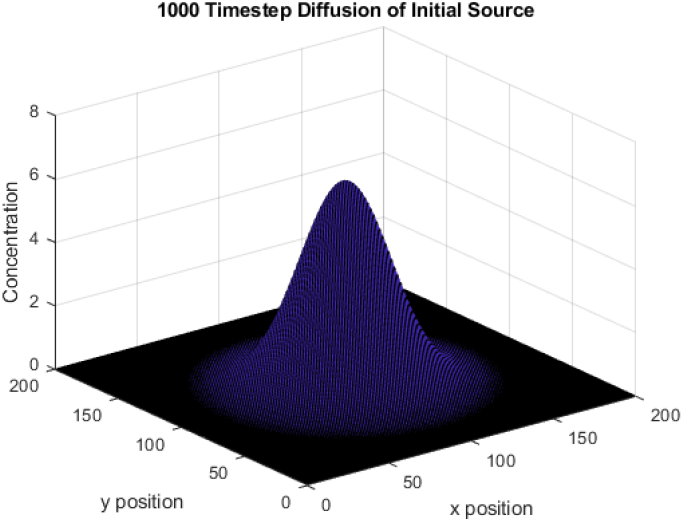
System Validation. Diffusion from a point source accurately produces a Gaussian concentration profile.

## RESULTS AND DISCUSSION

### Animal Morphology

How do animals get their shapes? Although dozens of signaling pathways from Fgf to Bmp are involved [1], Alan Turing proposed that complex pattern formation can be described by a reaction diffusion system with only two factors and minor random fluctuations [2]. Turing solved this system by linearizing around the steady state, which prevents understanding of system behavior over long or divergent timeframes.

To model this numerically, the following system of two dummy morphogens, A and B, was used. The change in concentrations over time is affected by presence of the other factor, as well as a linear offset (Equations 4 and 5).

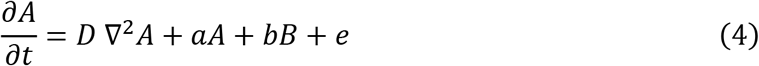

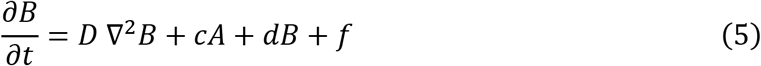

The system is initialized under some arbitrary constraints (Equations 6 and 7), including an arbitrary perturbation *ϵ*.

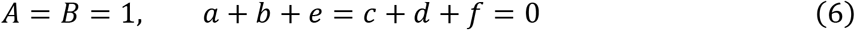

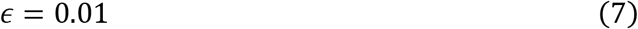

To break symmetry, we randomly generate pairs of fluctuations (these represent elements being thermally bounced “back and forth”), where the first element in the pair has concentrations A_1_, B_1_, and the second has concentrations A_2_, B_2_ (Equation 8).

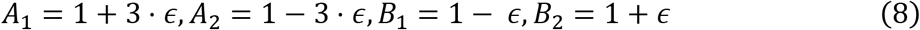

By varying *a - d* and altering the locations of pairs relative to each other, many morphologies are attainable.

Varying the values *a - d* results in changes in the density of heterogenous defects/lines in the steady state – higher values of coefficients result in more dense patterns (Figure 2). This means the difference between an animal with very complicated patterns everywhere and one with larger spots/defects may be faster reaction rates of morphogens in the more complex animal.

**Figure 2:**
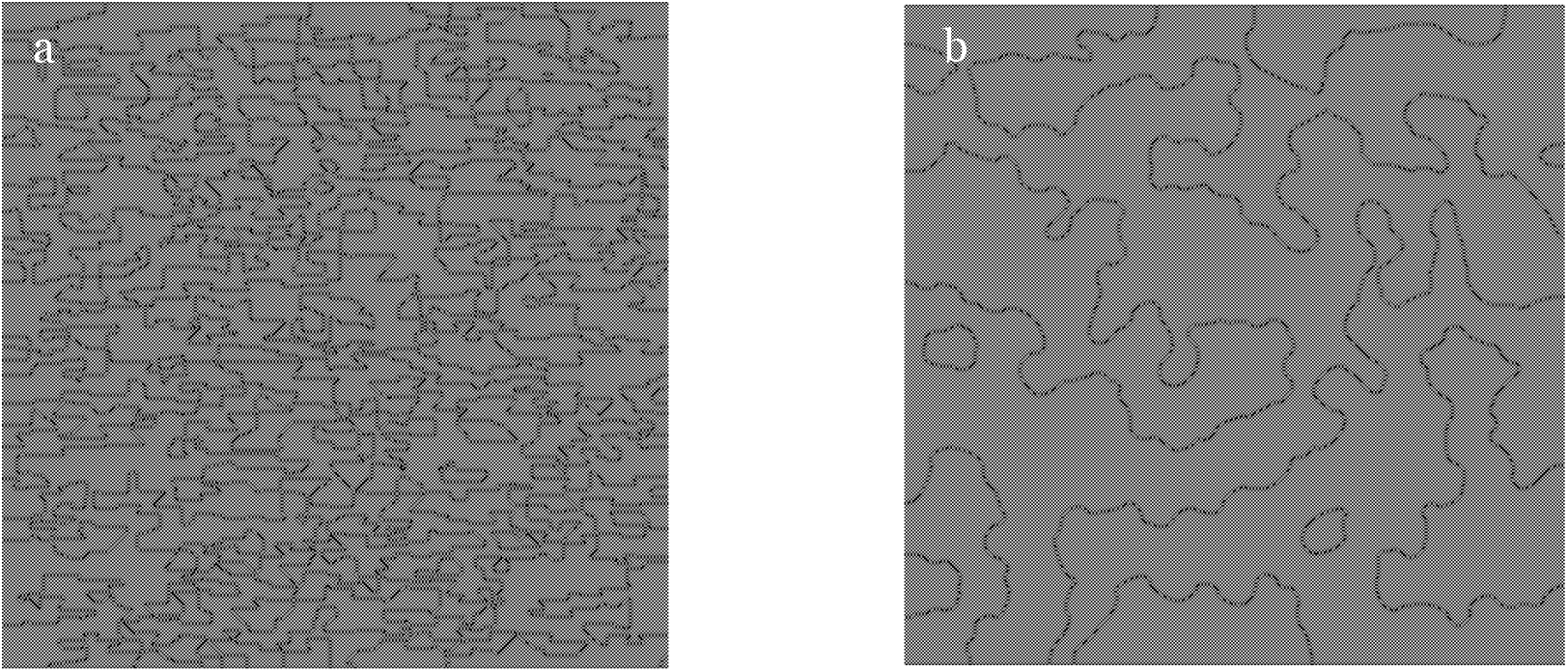
Representative Steady States from systems with 1000 randomly located point pairs. (a) Parameters: a = 100000, b = −100001, c = 100001, d = −1000002. (b) Parameters: a = 5, b = −6, c = 6, and d = 7.

Pair location has little effect – if each point is randomly located (i.e. the A1 defect is not next to the A2 defect), the same patterns are produced as when each point in the pair is one apart

This model helps demonstrate the failure of the linearization approximation (i.e. long-term stability even with predicted short-term non-equilibrium perturbations). Turing’s approximation is useful for some patterns, but over time some smaller aspects of these patterns disappear – this is why it is useful to numerically model the full timescale of pattern formation in order to observe the ultimate behavior at equilibrium (Figure 3).

**Figure 3:**
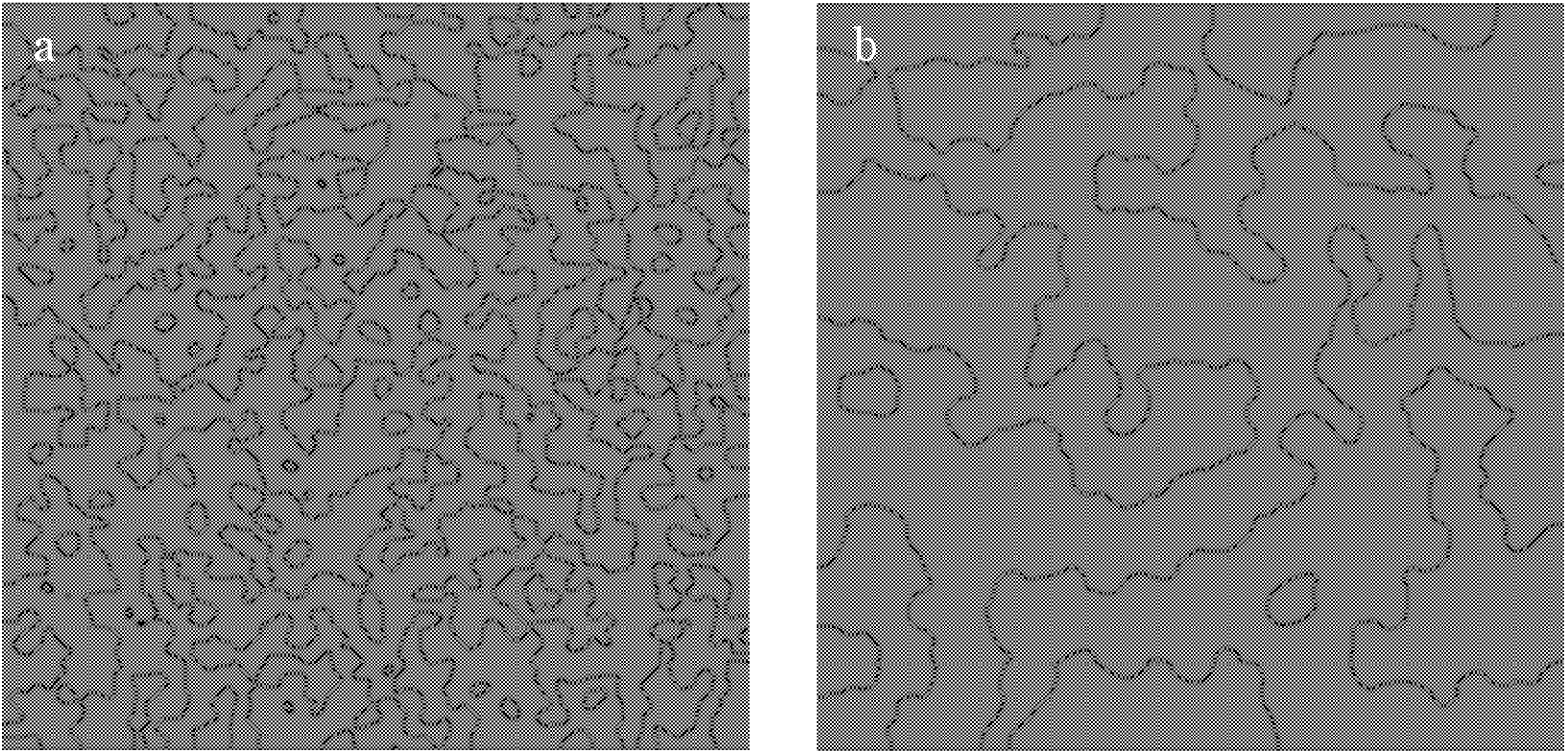
Time Dependence. (a) Short-term linear approximation of diffusing factors, as predicted by Turing’s model. (b) True, long-term steady-state.

These pattern formations are more than just mathematical curiosities. By varying the number of random defects, constants, and locations of initial defects (as organisms do during development), patterns resembling giraffe spots and even drosophila embryo segmentation appear (Figure 4).

**Figure 4:**
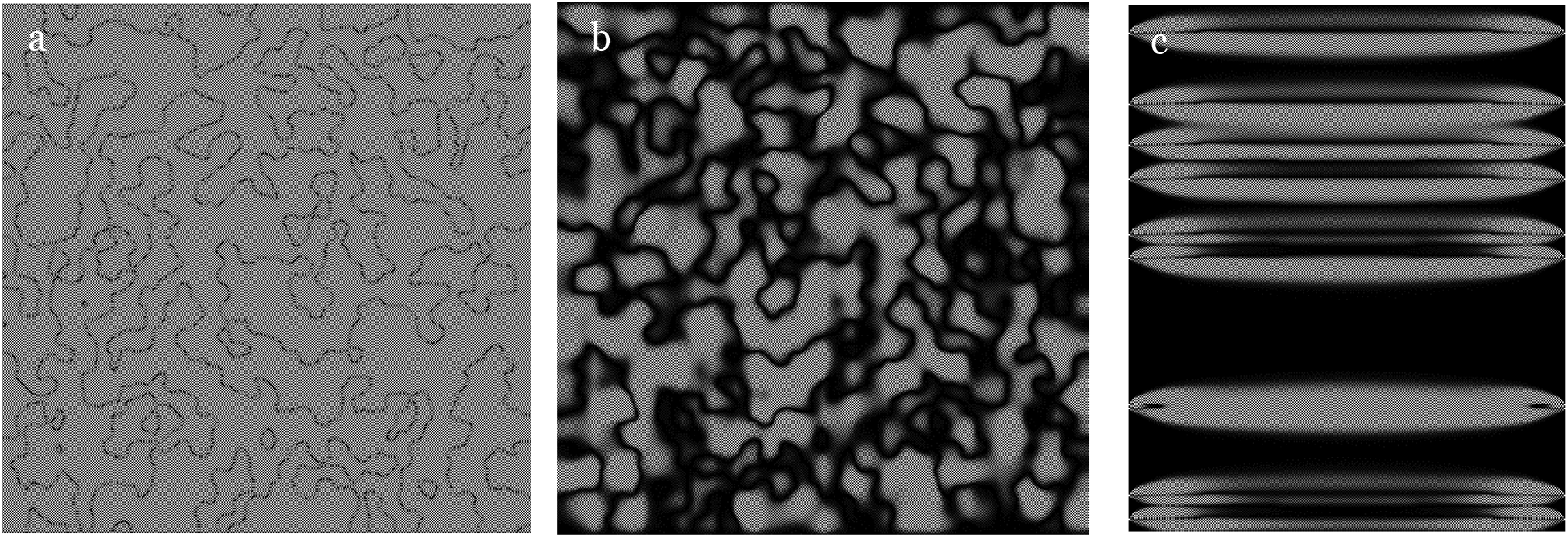
Animal Morphologies. (a-b) “Giraffe Spots.” (c) “Drosophila Segments.”

### Modeling Bacteria in Non-Equilibrium Growth

While reaction diffusion on its own leads to interesting conclusions, it is instructive to incorporate growth into the model, given that cells do not exist in a static environment and divide under favorable conditions. Ben-Jacob observed interesting growth regimes when growing bacteria on agar with different peptone concentrations [3, 4]. Using the numerical model, the real patterns of Figure 5 can be modeled mathematically using reasonable physical constants.

**Figure 5:**
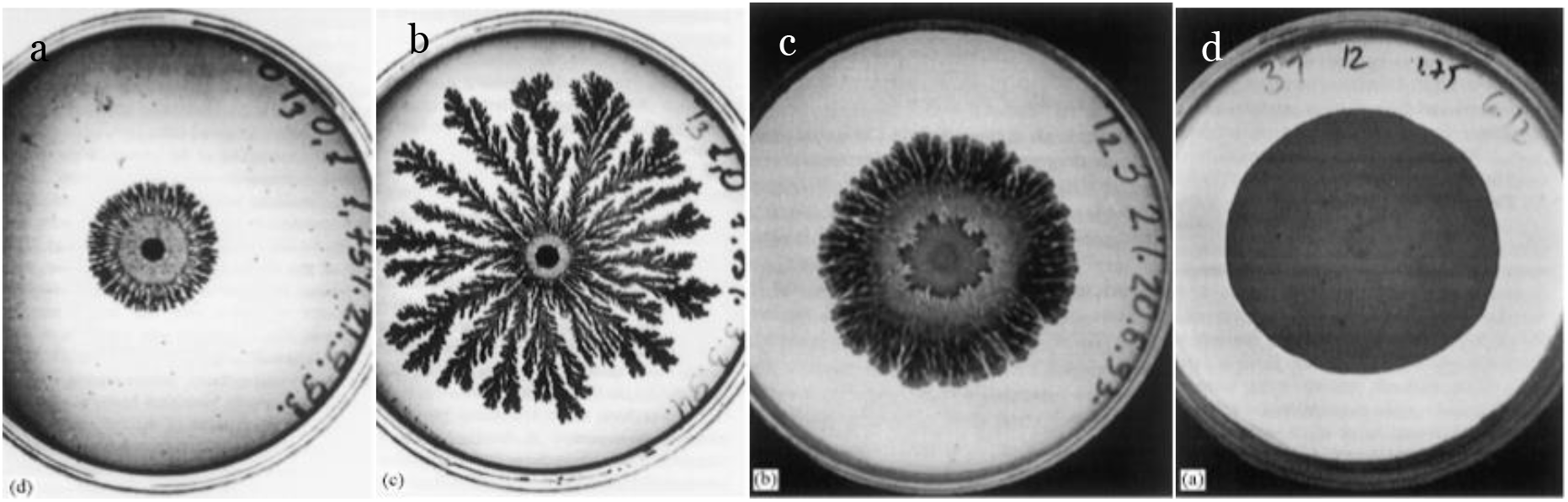
Empirical bacteria patterns [4]. Different growth regimes result from varying peptone concentrations of 0.1, 1, 3, and 10 g/l, respectively (a-d).

To model bacterial growth, assume each pixel represents one bacterium. Since a bacterial radius is about 5 microns, a pixel is approximated as a single 10×10 micron bacteria. Each bacterium has a certain uptake rate of nutrients, and will divide when the nutrient concentration exceeds a threshold; each resulting bacterium will have half the nutrients of the initial bacteria. In this model, there is no diffusion of bacteria themselves.

In addition, note that the expected length of diffusion is described by the following equation:

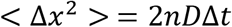

For reasonable physical constants, set D = 10^−7^ cm^2^/s and Δ*x* to the size of a bacterium (10 microns), yielding a timestep of 5 seconds. Additionally, let the amount of nutrients a bacterium needs before division be 3E-12 grams. Finally, test a range of peptone concentrations from 1E-6 to 2E-6 g/cm^2^.

The numerical model reproduces the four growth regimes observed by Ben-Jacob’s group (Figure 6). At low peptone concentration, there is a circle in the center, and small radial branching. At intermediate peptone concentration, there is radial branching and fractal-like growth on these branches. At high peptone concentration, there is radial “finger” growth, and at very high peptone concentrations, a uniform bacterial blob grows. This suggests that bacterial growth is diffusion limited and can be modeled as such, in close agreement with reported experimental data.

**Figure 6:**
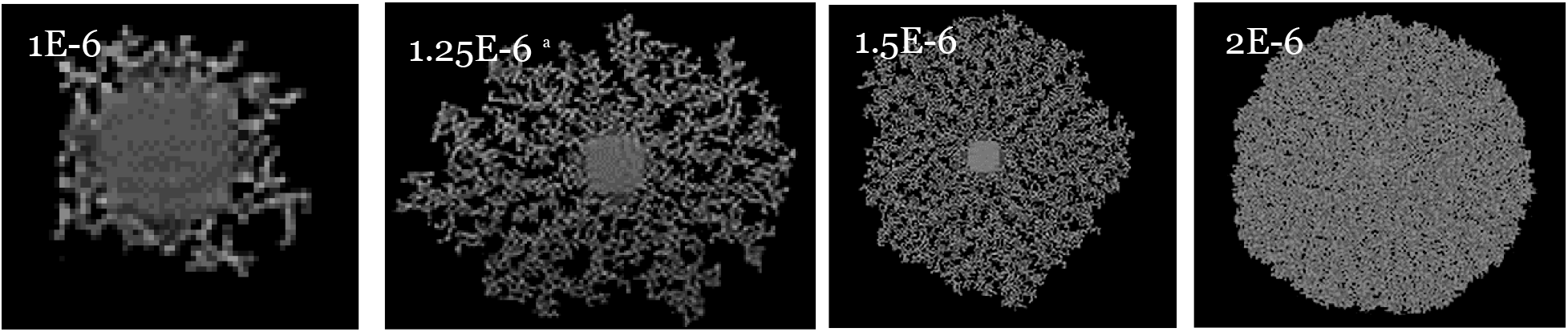
Modeled bacteria patterns. Labeled by peptone concentration (g/cm^2^).

## CONCLUSION

Complex systems of development or signaling pathways can be explained and modeled to a high degree of accuracy with a few simplifying assumptions, as demonstrated by the test cases of leopard spot morphogenesis and Ben-Jacob bacterial growth. As a result, diffusion-limited models have the potential to drive and explain advancements in developmental biology. Beyond biological applications, a numerical model of diffusion has applications to physical systems from fluid flow to heat transfer.

